# More than DNA methylation: does pleiotropy drive the complex pattern of evolution of *Dnmt1*?

**DOI:** 10.1101/824052

**Authors:** Ashley U. Amukamara, Joshua T. Washington, Zachary Sanchez, Elizabeth C. McKinney, Allen J. Moore, Robert J. Schmitz, Patricia J. Moore

## Abstract

DNA methylation is an important chromatin modification that can stably alter gene expression in cells and maintain genome integrity in plants and vertebrates. The function of DNA methylation outside of these well-studied systems, however, is unclear. Insects, in particular, represent an understudied group. Variation in the level of DNA methylation and gains and losses in the maintenance methyltransferase, DNMT1, across the insect tree of life suggests that there is much we don’t understand about DMNT1 function and evolution. One constant across the studies examining patterns of *Dnmt1* expression in insects is that expression is consistently high in reproductive tissues compared to somatic tissue. The explanation for this has been that DNMT1 is required in tissues that have high levels of cell division. Our previous study found that downregulation of *Dnmt1* expression in the milkweed bug *Oncopeltus fasciatus* results in the expected reduction of DNA methylation, no global changes in gene expression reflecting changes in DNA methylation, and the loss of the ability to produce viable oocytes. Here, we show that females treated with ds-*Dnmt1* RNA during larval development have a more extreme phenotype; they lack oocytes entirely but develop a normal somatic ovary. Our results indicate a specific role for DNMT1 in the formation of gametes and are consistent with data from other systems, including *Tribolium castaneum*, a species does not have DNA methylation. We propose that DNMT1 has multiple functional roles in addition to methylating DNA, which explains its complex patterns of evolution, and suggests that previous inferences of causation from associations are premature.

## Introduction

DNA methylation is an important chromatin modification that can stably alter gene expression in cells and it functions in some species in the maintenance of genome integrity (Law and Jacobson 2010, Schmitz et al. 2019). Although diverse roles for DNA methylation have been identified in some plant, animal, and fungal species (Schmitz et al. 2019), the function of DNA methylation in animals outside of well-studied vertebrate systems is unclear. In particular, DNA methylation in the insects has been understudied. This is unfortunate, as there is extensive variation in levels of DNA methylation, from no methylation to high methylation, across the insect tree of life (Bewick et al. 2016, Glastad et al. 2019). Variation exists in the amount of methylated DNA as well as the enzymatic toolkit required to establish and maintain it (Bewick et al. 2016). DNA methylation is associated with a suite of DNA methyltransferases (DNMT; Lyko 2017), which is generally found across eukaryotic genomes (Schmitz et al. 2019). The canonical DNMT enzymes include DNMT1, which maintains existing patterns of symmetrical DNA methylation by restoring the opposing methyl group to a cytosine residue in a hemimethylated CpG following DNA replication and DNMT3, which can methylate cytosines *de novo*. Insects even vary in these typically conserved genes. The phylogenetic distribution of DNMT1 in insects is more widespread than DNMT3 and is duplicated in some groups. Curiously, DNMT1 is found in insect taxa with low or no DNA methylation, including the red flour beetle *Tribolium castaneum* (Schulz et al. 2018). This variation raises questions both about the function of DNA methylation in insects and the selection leading to differences among taxa.

In this study we have investigated the function of DNMT1 in the large milkweed bug, *Oncopeltus fasciatus*, an evolution and development model system with molecular genetic resources, a methylated genome and the full complement of DNA methyltransferases. Using parental RNAi, we have previously shown that knockdown of *Dnmt1* in sexually mature females results in embryo arrest (Bewick et al. 2019). Eventually, females injected with ds-*Dnmt1* stopped laying eggs and their ovarian structure was significantly disrupted. We further found that reduction of DNA methylation was not directly associated with changes to gene expression patterns within the ovary and that DNA methylation patterns in the somatic tissues, the gut and muscles, were unaffected by the knockdown. Our parental RNAi experiments raise the question as to why the morphological defects of *Dnmt1* knockdown are confined to the ovary if the role of *Dnmt1* is to maintain a methylated genome in all cells.

The most parsimonious hypothesis for why only the ovary displays a phenotype following *Dnmt1* knockdown is that there are not enough rounds of DNA replication in the somatic cells between injection of the ds-*Dnmt1* into the adult and sampling of the tissues for any phenotype to be realized. An alternative hypothesis is that there is an additional role for *Dnmt1* in reproductive tissue. This latter hypothesis is partially supported by expression patterns of *Dnmt1* in other insects. A common feature of *Dnmt1* and *Dnmt3* expression patterns in several species of insects is that it is often correlated with reproductive tissues (Kay et al. 2018), leading to the hypothesis that these enzymes may play a role in gametogenesis, gamete quality, or early embryogenesis. While only a few studies have examined the functional role of *Dnmt1* knockdown on DNA methylation levels in insects, similar morphological phenotypes have been observed in other insects following *Dnmt1* knockdown. In the jewel wasp *Nasonia vitripennis*, there are three distinct *Dnmt1* genes, all of which are expressed and maternally provided to the embryo. One of these, *Dnmt1a* is required for early embryogenesis (Zwier et al. 2012). *Tribolium castaneum*, which lacks a methylated genome but retains *Dnmt1*, also has increased expression of *Dnmt1* in female reproductive tissue. While *Dnmt1* is expressed at all life stages, it is upregulated in ovaries, suggesting a role in reproduction. Finally, parental RNAi of *Dnmt1* demonstrated that *T. castaneum* requires maternal expression of *Dnmt1* for early embryogenesis (Schulz et al. 2018). Together, these studies suggest a germ cell specific function for *Dnmt1* in insects.

In this paper, we examine whether there is a specific role for *Dnmt1* in the development of germ cells outside of its function in maintaining DNA methylation. To differentiate among our two alternative hypotheses, we injected individuals during larval development to allow for sufficient time and rounds of cell division for DNA methylation to be lost in somatic cell genomes and for any somatic morphological phenotype to be manifest. We could reject the insufficient rounds of cell division explanation, so we then tested if *Dnmt1* is required for oogenesis, but not required for somatic cell function.

In the first set of experiments, we injected fourth instar nymphs to provide adequate rounds of cell division between downregulation of *Dnmt1* expression and adult stages. *O. fasciatus* is a hemimetabolous insect, undergoing incomplete metamorphosis. It develops through five immature instar stages; at each stage there is further development of color, wings and genitalia. *O. fasciatus* have telotrophic, meroistic ovaries with seven ovarioles per ovary. Female first instar nymphs hatch with primordial ovaries in which the female germ cells, the oogonia, have been set aside during embryogenesis (Wick and Bonhag 1955). During the second and third instar stages, the ovaries increase in size and there is an increase in the numbers of oogonia by mitotic cell division. During the fourth instar stage, some oogonia divide mitotically to form trophocytes while others divide meiotically to form gametes, the primary oocytes. The trophic tissues (equivalent to nurse cells) differentiate during the fifth and final instar stage and at adult emergence, the ovary consists of nutritive trophic tissue, oogonia, primary oocytes, small growing oocytes connected to the nutritive tissue by trophic cords, and prefollicular cells that will envelop the developing oocytes. As the female undergoes sexual maturation, the oocytes move into the ovariole and complete maturation.

Our results established that knockdown of *Dnmt1* earlier in development did not result in somatic morphological phenotypes, but rather exacerbated the impact on germ cell development. Given these results, we focused our experiments on testing the hypothesis that *Dnmt1* is required for oogenesis, but is not required for somatic cell function. As a control, we included *Boule*, a gene known to be required for oogenesis in *O. fasciatus* (Ewen-Campen et al. 2013) for comparison. Downregulation of *Boule* expression in adult females using RNAi results in an almost identical phenotype to that of downregulating *Dnmt1* (Ewen-Campen et al. 2013); initially, sexually mature females injected with ds-*Boule* lay eggs that don’t develop, but after a few clutches, females stop producing eggs. *Boule* is a widely conserved gene required for reproduction across the bilateral animals (Shah et al. 2010). Mutations in *Boule* cause arrest in meiosis prior to metaphase in *Drosophila melanogaster* males and *Caenorhabditis elegans* females (Karashima et al. 2000). Thus, *Boule* provided an excellent comparison for evaluating the hypothesis that *Dnmt1* is required for oogenesis, and perhaps the transition from oogonia to oocytes in *O. fasciatus*. We examined this hypothesis by testing for the following predictions arising from our overall hypothesis. First, *Dnmt1* should be most highly expressed when and where oogenesis is occurring. Second, downregulating *Dnmt1* expression during a critical developmental stage for oogenesis would result in the loss of oocyte production. Third, downregulating *Dnmt1* expression will affect reproductive function without affecting somatic function and lifespan.

## Methods

### Animal husbandry

To produce nymphs of known age and social history, *O. fasciatus* adults from our mass colony were randomly mated, and resulting eggs were removed and stored in individual plastic containers. Upon hatching, nymphs were transferred into 2-cup Rubbermaid containers supplied with organic, raw sunflower seeds and deionized water. Nymph colonies were housed under 12 hr:12 hr light/dark conditions at 27°C. Nymphs were staged utilizing a visual development chart from Chesebro et al. (2009).

### Nymph RNAi treatment

DNA templates of *Dnmt1* and *Boule* was prepared using a PCR reaction with gene-specific primers (Table 1). Double stranded RNA was synthesized with an Ambion MEGAscript kit (ThermoFisher Sci, Waltham, MA) to generate sense and anti-sense RNA which were allowed to anneal. The concentration of ds-RNA was adjusted to 4 µg/µL in injection buffer (5 mM KCl, 0.1 mM NaH_2_PO_4_) for both *Dnmt1* and *Boule*. Female fourth instar nymphs, hatched within 2-3 days of each other, were selected for injection. Treatments were ds-*Dnmt1*, ds-*Boule*, and control injections with injection buffer alone. Nymphs were injected with 2 µL using a syringe and a pulled glass capillary needle (Sutter Instrument Company micropipette puller model P-97, Novato, CA) between the third and fourth abdominal segments (Chesebro et al. 2009). Injected nymphs were housed in petri dishes with *ad libitum* sunflower seeds and water under standard rearing conditions until they emerged as adults.

**Table 1.**
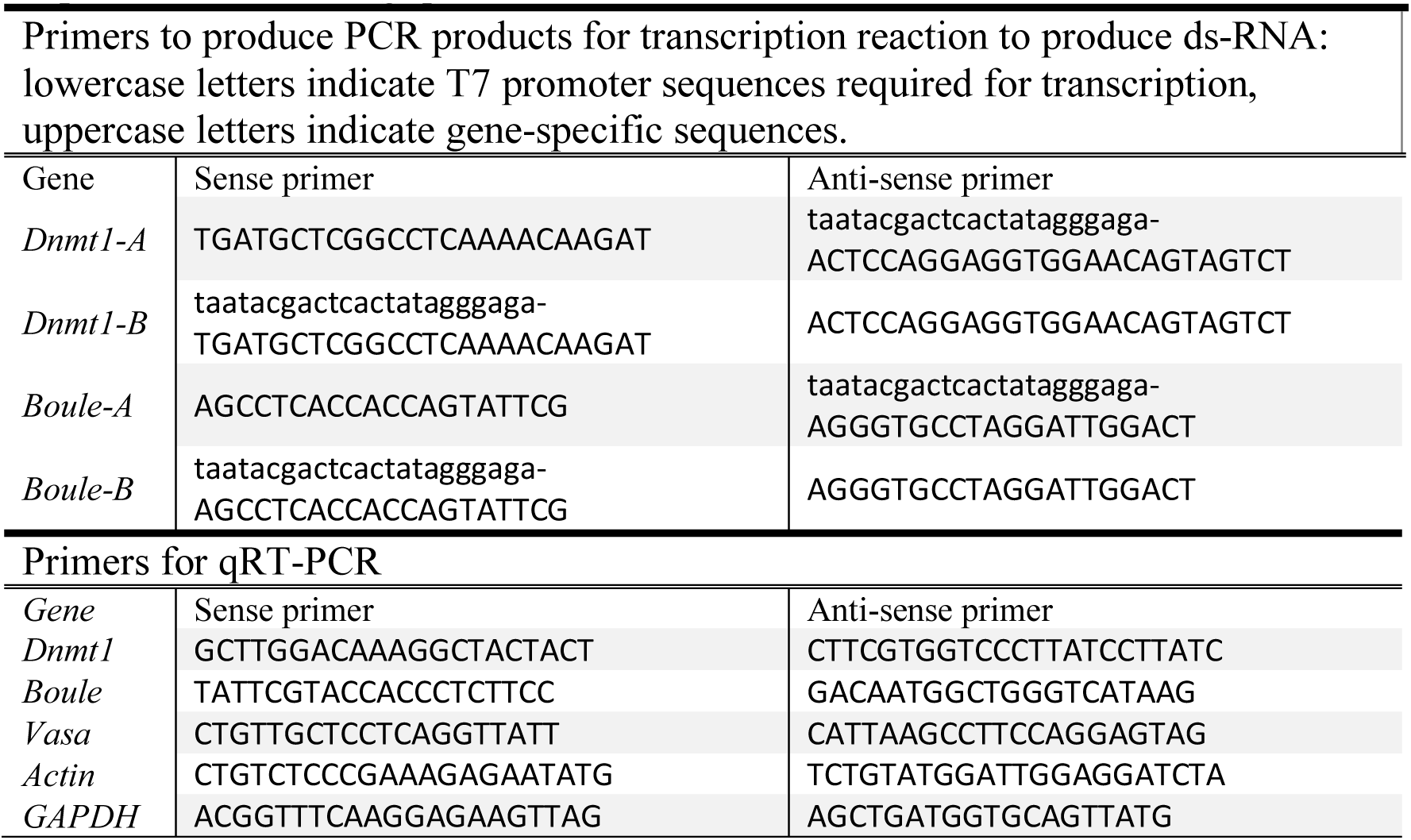
Primer sequences used to generate RNAi for injection and to quantify expression levels using quantitative Real Time PCR

### qRT-PCR, development series, and tissue specificity

To confirm that our RNAi treatment during nymphal development effectively knocked down expression of *Dnmt1* in 21-day old adults emerging from treated nymphs, total RNA was extracted from whole bodies of flash frozen adults from either control or ds-*Dnmt1* treated individuals using a Qiagen RNA Easy kit with Qiazol (Qiagen, Venlo, The Netherlands) as per manufacturer’s instructions. Complementary DNA (cDNA) was synthesized from 500 ng RNA with qScript cDNA Super-Mix (Quanta Biosciences, Gaithersburg, MD). Expression levels of *Dnmt1* were quantified by quantitative real-time PCR (qRT-PCR). Primers were designed using the *O. fasciatus* genome as a reference (Panfilio et al. 2019) and actin and GAPDH were used as endogenous reference genes (Table 1). We used a Roche LightCycler 480 with the SYBR Green Master Mix (Roche Applied Science Indianapolis, IN). All samples were run with 3 technical replicates using 10 μL reactions. Primer efficiency calculations, genomic contamination testing, and endogenous control gene selection were performed as described in Cunningham et al. 2014. We used the ΔΔCT method to compare levels of gene expression across the samples (Livak and Schmittgen 2001). *Dnmt1* expression is significantly knocked down in adults injected with ds-*Dnmt1* as fourth instar nymphs (ANOVA; F = 19.038, d.f. = 1, 18, p < 0.001).

To examine expression levels of *Dnmt1* across development, groups of nymphs were staged and flash frozen in liquid nitrogen, and stored at −80°C. The earliest stages of nymphs are difficult to sex accurately, so the 2^nd^ and 3^rd^ instar nymphs were not separated by sex. By the 4^th^ instar stage of development, females can be identified, so only female 4^th^ and 5^th^ instar nymphs were collected. To ensure that there was a sufficient amount of tissue for RNA isolation, smaller nymphs were pooled for each sample. A sample of 2^nd^, 3^rd^, 4^th^ and 5^th^ instar nymphs contained 10, 5, 4, and 3 individuals, respectively. The other two developmental stages we collected were newly emerged females, collected within 24 hours of adult emergence, and sexually mature females, collected 7-10 days post-adult emergence. All adult females were housed in a container without males to ensure that they had not mated prior to RNA isolation. A sample of adults consisted of a single individual. Total RNA was extracted and cDNA was synthesized as described above.

Expression levels of *Dnmt1, Boule* and *Vasa* were quantified by quantitative real-time PCR (qRT-PCR). *Boule* and *Vasa* primers were designed using the *O. fasciatus* genome as a reference (Panfilio et al. 2019) and actin and GAPDH were used as endogenous reference genes (Table 1). Expression levels were quantified by qRT-PCR as described above. We used ANOVA followed by Tukey-Kramer HSD to compare all pairs in JMP Pro 14.1 to determine significant differences in levels of gene expression among the different stages.

To examine for tissue specific expression of *Dnmt1*, we dissected adult, virgin females 7-10 days post adult emergence. We flash froze ovaries, gut, and thorax (muscle) from individual females. Total RNA was extracted and complementary DNA (cDNA) was synthesized as described above. Expression of *Dnmt1* was quantified and analyzed as described for the development series.

### Quantification of DNA methylation

To determine the effect of downregulation of *Dnmt1* expression from the fourth instar stage, total DNA was extracted from gut, muscle, and ovarian tissue samples from adult females that had been treated with ds-*Dnmt1* or control injections at the fourth instar using a Qiagen Allprep DNA/RNA Mini Kit (Qiagen, Venlo, The Netherlands). MethylC-seq libraries were prepared from the DNA isolated from the gut, muscle, and ovary tissue from females as described (Ulrich et al. 2015). Briefly, genomic DNA was sonicated to 200 bp using a Covaris S-series focused ultrasonicator and end-repaired using an End-It DNA end-repair kit (Epicentre, Madison, WI). The end-repaired DNA was subjected to A-tailing using Klenow 3′-5′ exo− (New England Biolabs, Ipswich, MA) and ligated to methylated adapters using T4 DNA ligase (New England Biolabs, Ipswich, MA). The ligated DNA was subsequently bisulfite converted using an EZ DNA Methylation-Gold (Zymo Research, Irvine, CA) kit as per the manufacturer’s instructions and amplified using KAPA HiFi Uracil + Readymix Polymerase (ThermoFisher Sci, Waltham, MA). Samples were sequenced on a NextSeq500 and qualified reads were aligned to the *O. fasciatus* genome assembly 1.0 as described (Schmitz et al. 2013). Chloroplast DNA (which is fully unmethylated) was used as a control to calculate the sodium bisulfite reaction non-conversion rate of unmodified cytosines (Supplementary Table 1).

### Fertility and fecundity assays

After the injected nymphs eclosed, the females were placed in individual petri dishes with *ad libitum* food and water and provided with loose cotton wool as an oviposition substrate. Injected females were paired with stock males from the mass colony for mating. Eggs were collected three times a week over a period of 2 weeks. Eggs were incubated in solo plastic cups at 27°C for one week to allow for embryonic development and hatching. The total numbers of eggs and the number of eggs that were fertilized and developed were counted. *Oncopeltus fasciatus* eggs are pale yellow in color when they are first laid. Fertilized eggs turn a deep orange as development proceeds, while unfertilized eggs remain pale yellow.

### Ovarian structure analysis

A separate set of female nymphs injected with either control, *Dnmt1*, or *Boule* ds-RNA was used to analyze the effect of knockdown on ovarian structure. Virgin females were dissected 7-10 days post-adult emergence. At this age, females are sexually receptive and have maturing oocytes in their ovaries. Females were dissected and the ovaries removed. For some females, whole mounts of ovaries were imaged using Leica M60 stereomicroscope and LAS v4 imaging software. All ovaries were processed for microscopy using the technique described in Duxbury et al. 2017. Briefly, ovaries were fixed for 30 minutes in 4% formaldehyde in Phosphate Buffered Saline plus 0.1% Triton-X100 (PBT) and stained for evidence of cells in the M phase of the cell cycle using an α-pHH3 primary antibody (Millipore antibody 06-570, Sigma-Aldrich, St. Louis, MO). The secondary antibody was an Alexa Fluor goat-anti-rabbit 647 (ThermoFisher Scientific, Waltham, MA). Following antibody staining the ovaries were stained with Invitrogen fluorescein labeled phalloidin (0.2 units/mL PBT; ThermoFisher Scientific, Waltham, MA) to visualize actin and DAPI (0.5 μg/mL PBT) to visualize nucleic acids. Stained ovaries were mounted in Mowiol 4-88 mounting medium (Sigma-Aldrich, St. Louis, MO) and imaged with a Zeiss LSM 710 Confocal Microscope (Zeiss) at the UGA Biomedical Microscopy core or an EVOS FI Cell Imaging system (ThermoFisher Scientific, Waltham, MA).

### Lifespan analysis

Control and experimental females from the fertility and fecundity assays were checked every 24 hours for viability. The date of adult emergence and the date of death were recorded for each female and lifespan was calculated as the number of days between adult emergence and death. We tested for differences in female longevity relative to treatment using a Wilcoxon Rank Sum test with JMP Pro V14.1.

## Results

Fourth instar nymphs injected with ds-*Dnmt1* are equally likely to survive to adulthood as control injected fourth instar nymphs. At least 80% of nymphs injected at the fourth instar stage emerged as adults in both control and ds-*Dnmt1* injection treatments. There was no difference in the likelihood of adult emergence among control and *Dnmt1* knockdown treatments (Contingency analysis: N = 80, d.f. = 1, Pearson χ^2^ = 0.082, p = 0.775). Furthermore, there was no obvious difference in morphology amongst the control and *Dnmt1* knockdown emerging adults. Adults from all treatments appeared normal, including apparently normal wing development. There were no visible phenotypic effects of reduced methylation in somatic tissue.

As predicted, injecting earlier in development resulted in reduced DNA methylation of all tissues due to an increased number of mitotic division cycles within the somatic tissues between treatment and sampling. In both somatic tissues we tested, gut and muscle, and in the ovary, methylated CpG levels went from around 12.5% in controls to around 5% in *Dnmt1* knockdowns (Figure 1).

**Figure 1.**
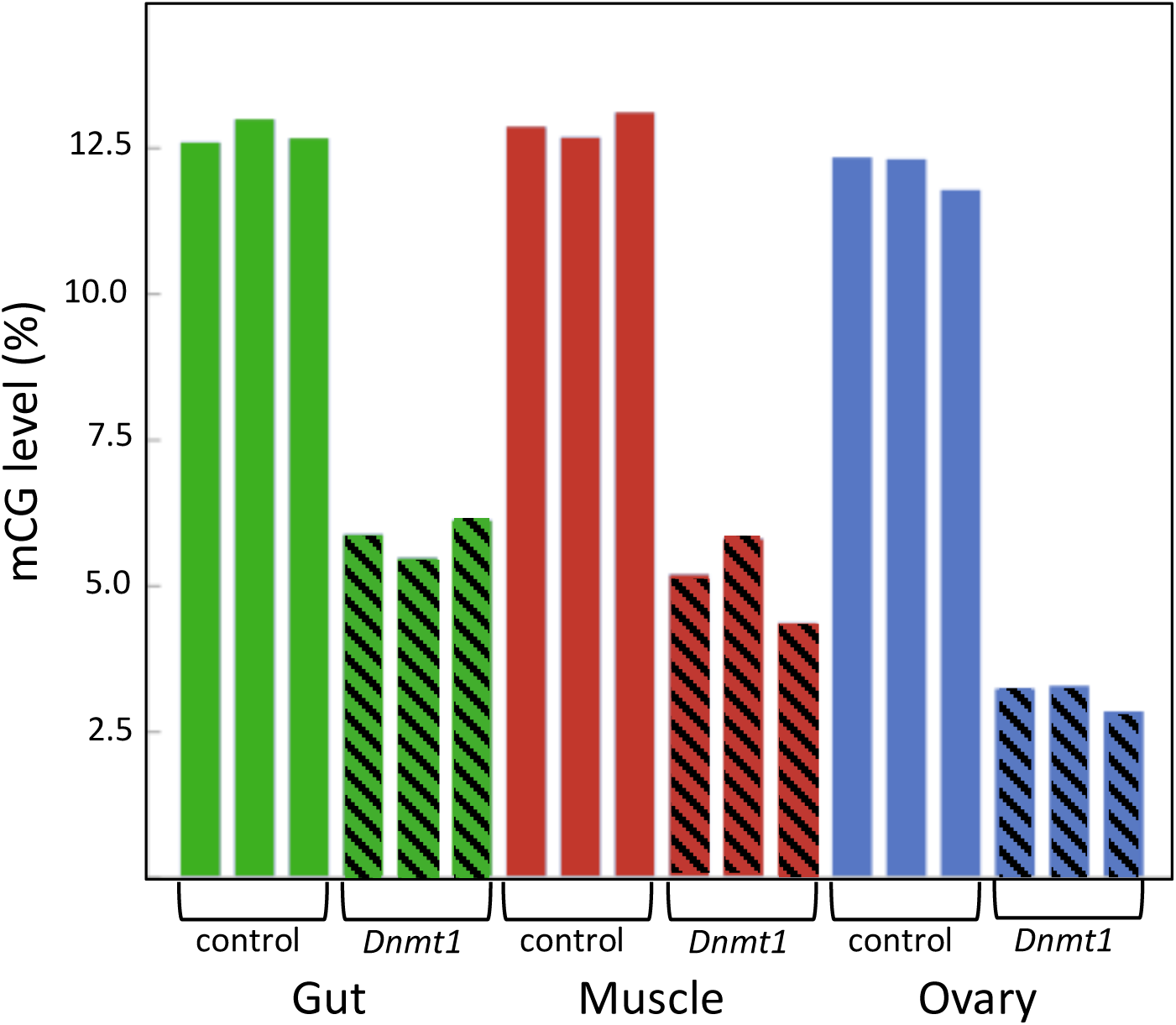
*Dnmt1* is required for the maintenance of mCG in somatic and reproductive tissue in *O. fasciatus*. Across all tissues the percent of mCG is consistently reduced in the genomes of individuals in which *Dnmt1* has been knocked down at the 4^th^ instar stage of development (hatched bars) compared to control individuals (solid bars).

### *Dnmt1* knockdown during ovarian development exacerbated impact on oogenesis

Examination of the ovaries from females emerging from fourth instar injected nymphs demonstrated a significant impact on the ovary: although the somatic ovary appeared to have developed normally, there were no eggs present (Figures 2A and 2B). *Dnmt1* knockdown clearly impacted female fertility. Only one out of 11 ds-*Dnmt1* treated female ever laid eggs across their entire lifespan, whereas all control females (N = 15) laid eggs. The number of eggs laid by this single ds-*Dnmt1* treated female was 86, whereas the control females laid a mean of 200 ± 42 eggs (Figure 2C).

**Figure 2.**
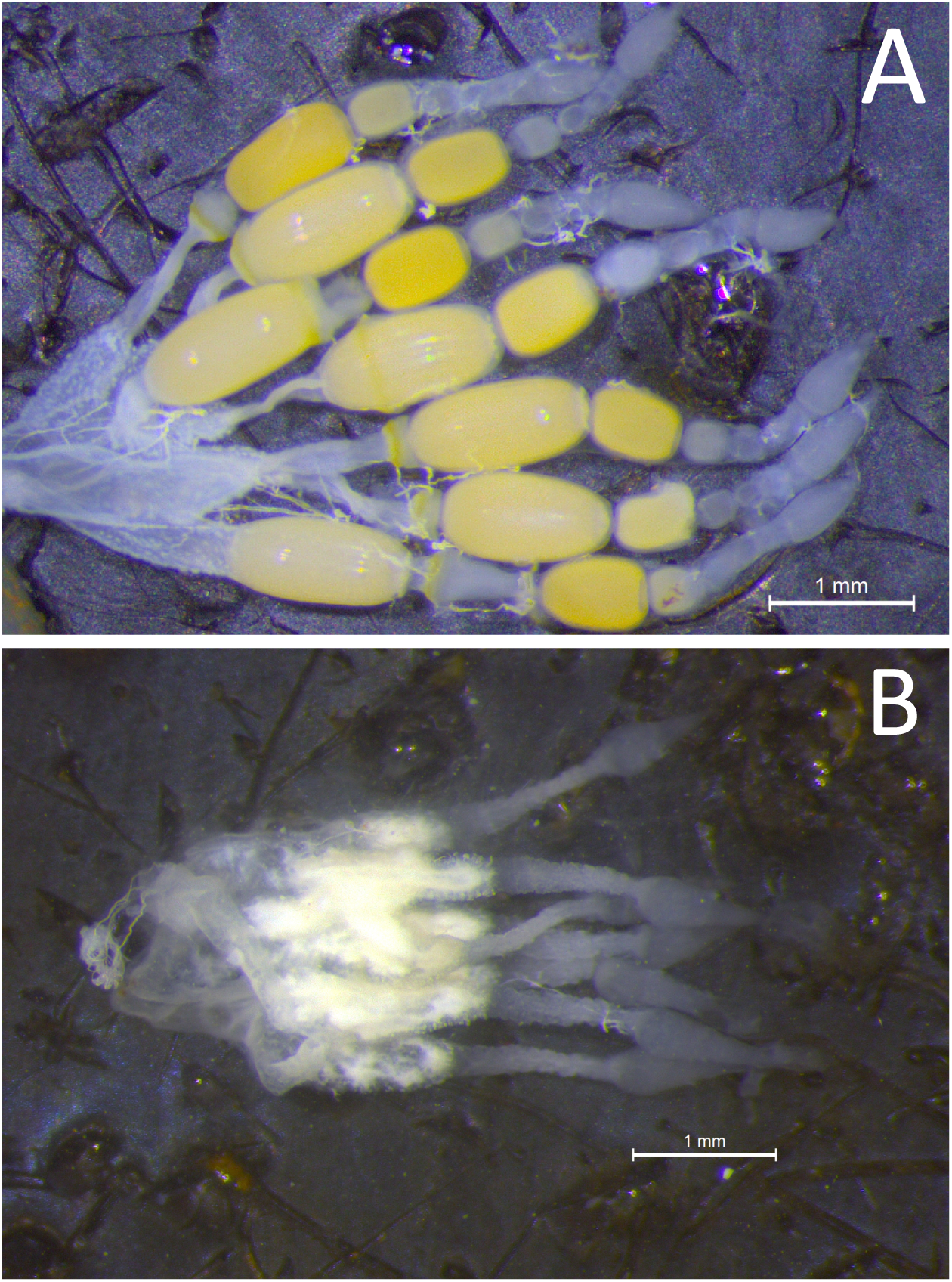
Whole mounts of ovaries from control (A) and *Dnmt1* knockdown (B) females demonstrated a significant impact on the ovary. While the somatic ovary appeared to have developed normally in the ds-*Dnmt1* females, there were no eggs present in the ovaries from adult females injected with ds-*Dnmt1* at the 4^th^ instar stage and these females never produced eggs over their lifespan.

### *Dnmt1* is expressed when and where oogenesis is occurring

*Dnmt1* expression went up in sexually mature females and expression pattern mirrored that of the two other oocyte specific genes we tested, *Vasa* and *Boule* (Figure 3). For all three genes, expression was statistically significantly affected by developmental stage, with expression being highest in the sexually mature females (*Dnmt1*, F_5,53_ = 89.749, p < 0.001; *Vasa*, F_5,53_ = 61.289, p < 0.001; *Boule*, F_5,53_ = 99.598, p < 0.001), with similar patterns in all three genes. *Dnmt1* is expressed in all tissues. However, expression levels significantly depend on tissue type (Figure 4; F = 51.661, d.f. = 2, 21, p < 0.001). Tukey-HSD pairwise comparison of the different tissues showed that expression is highest in the ovary, and did not differ between the two somatic tissues.

**Figure 3.**
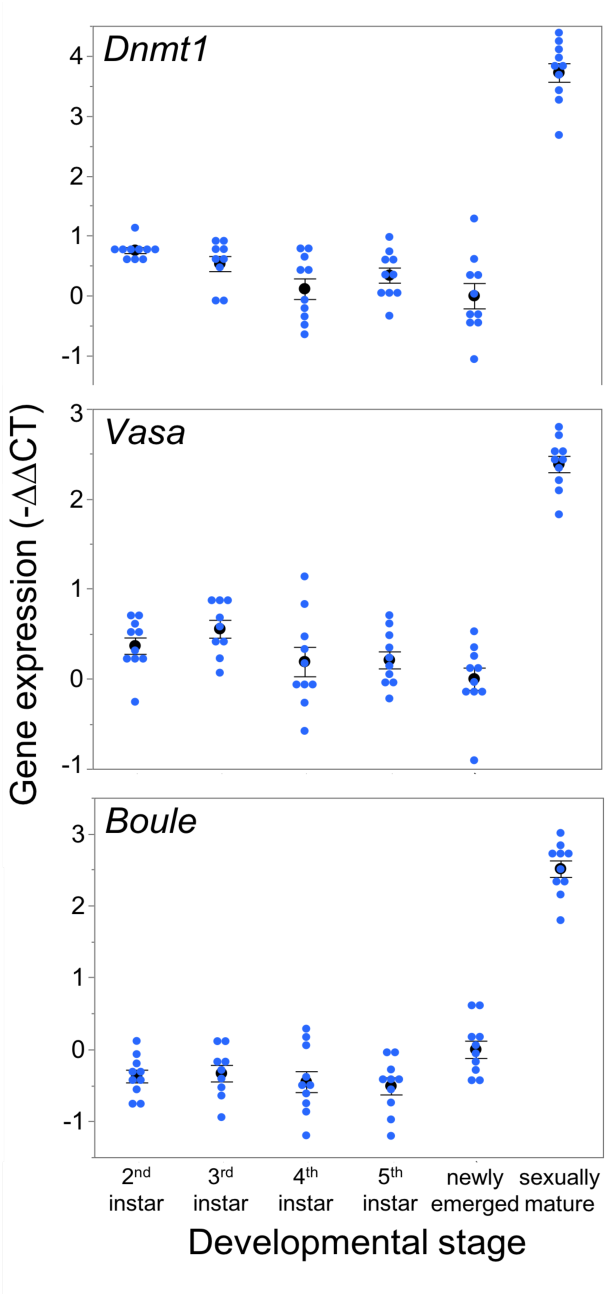
*Dnmt1* expression is highest in sexually mature females and the expression pattern of *Dnmt1* across development is the same as the germ cell specific genes *Vasa* and *Boule*. RNA was isolated from whole bodies of individuals at each instar stage of development from the 2^nd^ instar to the 5^th^ (final) instar. 2^nd^ and 3^rd^instar nymphs can not be sexed, but RNA was collected only from female 4^th^ and 5^th^ instar nymphs. Females were also sampled on the day of adult emergence and 10 days after adult emergence when they are sexually mature. Black dots and bars represent mean and SE. Blue dots represent data points for each individual female or nymph tested.

**Figure 4.**
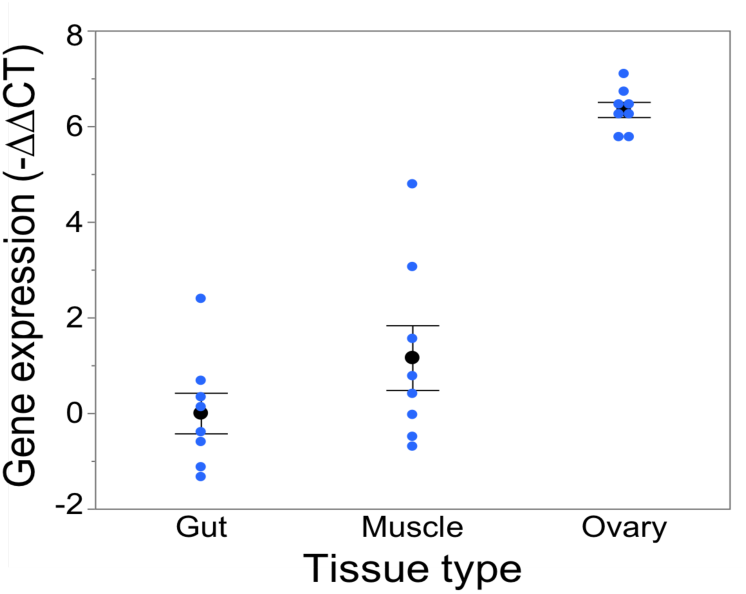
*Dnmt1* shows tissue specific expression. While *Dnmt1* is expressed in all tissues, it is more highly expressed in the ovary than in the two somatic tissues tested, gut and muscle. Letters designate significant differences among pairwise comparisons.

### *Dnmt1* knockdown leads to loss of developing oocytes, but not the supporting somatic cells

Closer examination of the structure of the ovaries from control, *Dnmt1* knockdown, and *Boule* knockdown females revealed that the somatic tissues within the ovary were all present in the ds-*Dnmt1* and ds-*Boule* treated females. The *O. fasciatus* ovary can be divided into a terminal filament, a germarium containing trophocytes, oogonia, primary oocytes, and prefollicular tissue, the vitellarium where oocytes mature, and the pedicel that connects the germarium to the oviduct and in which the developing oocytes mature (Figure 5A; see also Bonhag and Wick 1953). Low magnification images revealed that the somatic cells of the ovary, including the terminal filament, the tropic tissue, and the sheath cells that surround the developing ooctyes and extend to form the pedicel, were present in the ovaries of control females as well as *Dnmt1* females (Figures 5B-C). Trophic cells were easily recognized by their distinctive large nuclei in aggregates around the trophic core. *Dnmt1* knockdown females had trophic tissue with recognizable nuclear structure within the germarium region of the ovary and normal looking pedicels (Figure 5C). We also examined the structure of ovarioles from females in which *Boule* was knocked down from the fourth instar stage to compare the results from the *Dnmt1* knockdown females to those with a knockdown of a gene with a known function in gametogenesis (Ewen-Campen et al 2013). As with the ds-*Dnmt1* treated females, females treated with ds-*Boule* as fourth instar nymphs never formed oocytes, yet had recognizable trophic tissue in the germarium and normal pedicel structure (Figure 5D).

**Figure 5.**
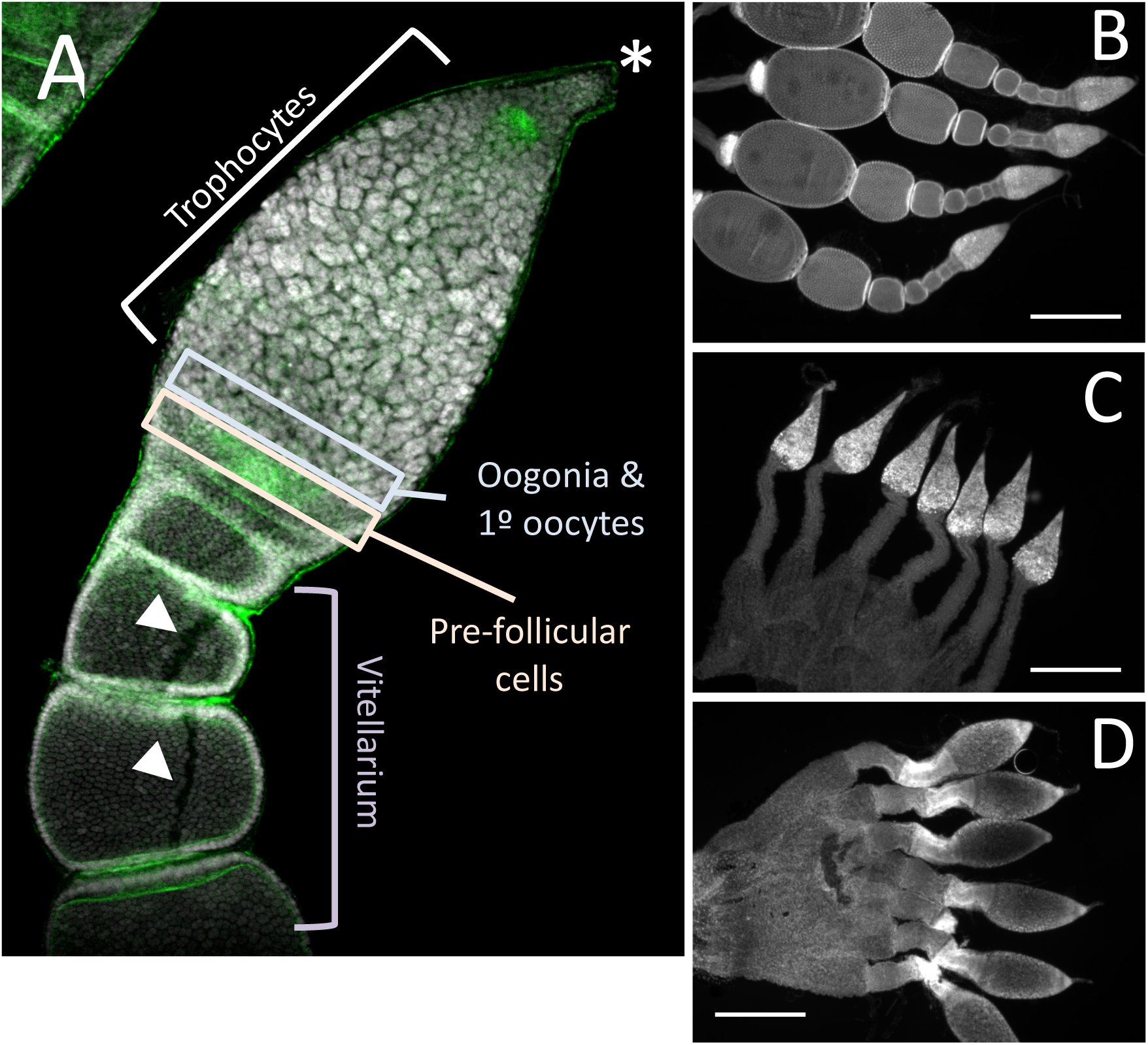
The structure of the somatic ovary, including the trophocytes, which develop from oogonia via mitotic division, was relatively normal in both Dnmt1 and Boule knockdown females. (A) Overview of the tip of an ovariole from a control female with the significant regions marked. The terminal tip (asterix) anchors the ovariole to the body wall. The majority of the germarium contains trophocytes that surround a trophic core. At the base of the germarium there is a group of cells with smaller nuclei. These cells are oogonia and primary ooctyes and pre-follicular cells that will envelope the developing oocyte as it moves into the vitellarium. During early stages of oocyte development, ooctyes remain connected to the trophocytes through trophic cords (arrowheads). Low magnification images (2X) of ovaries from control (B), *Dnmt1* knockdown (C), and *Boule* knockdown females stained with DAPI show that the germarium and the pedicel cells were present in the two knockdowns. However, no oocytes or their associated follicle cells ever entered the vitellarium.

Higher magnification of the germarium region of the ovarioles of control, ds-*Dnmt1*, and ds-*Boule* treated females (Figure 6) show that the large, aggregated nuclei of the trophocytes were present around a trophic core lined with actin in all three treatments. However, there are notable differences among the ds-*Dnmt1* treated females and the ds-*Boule* treated females in the region containing the oogonia and primary ooctyes. The germarium tissue in the *Boule* knockdown females remained organized and the different regions of cell types were recognizable. The small nuclei of the oogonia are apparent at the base of the germarium (Figure 6G). The germarium tissue of the *Dnmt1* knockdown females, however, was clearly disorganized and there was clear evidence of tissue disruption and cell death in the region where the oogonia and primary ooctyes would have been found (Figure 6D). Although the appearance of the control and ds-*Boule* germarium tissue was highly repeatable, there was significant variation in the amount of both cell death, evidenced by highly condensed and fragmented nuclei, and tissue disruption in the germarium of ds-*Dnmt1* treated females (Supplementary materials Figure S1). Despite the specific expression of the phenotype, all *Dnmt1* knockdown females exhibited defects in the same region of the germarium between the trophocytes and ovariole sheath cells where the oogonia and primary ooctyes would be located.

**Figure 6.**
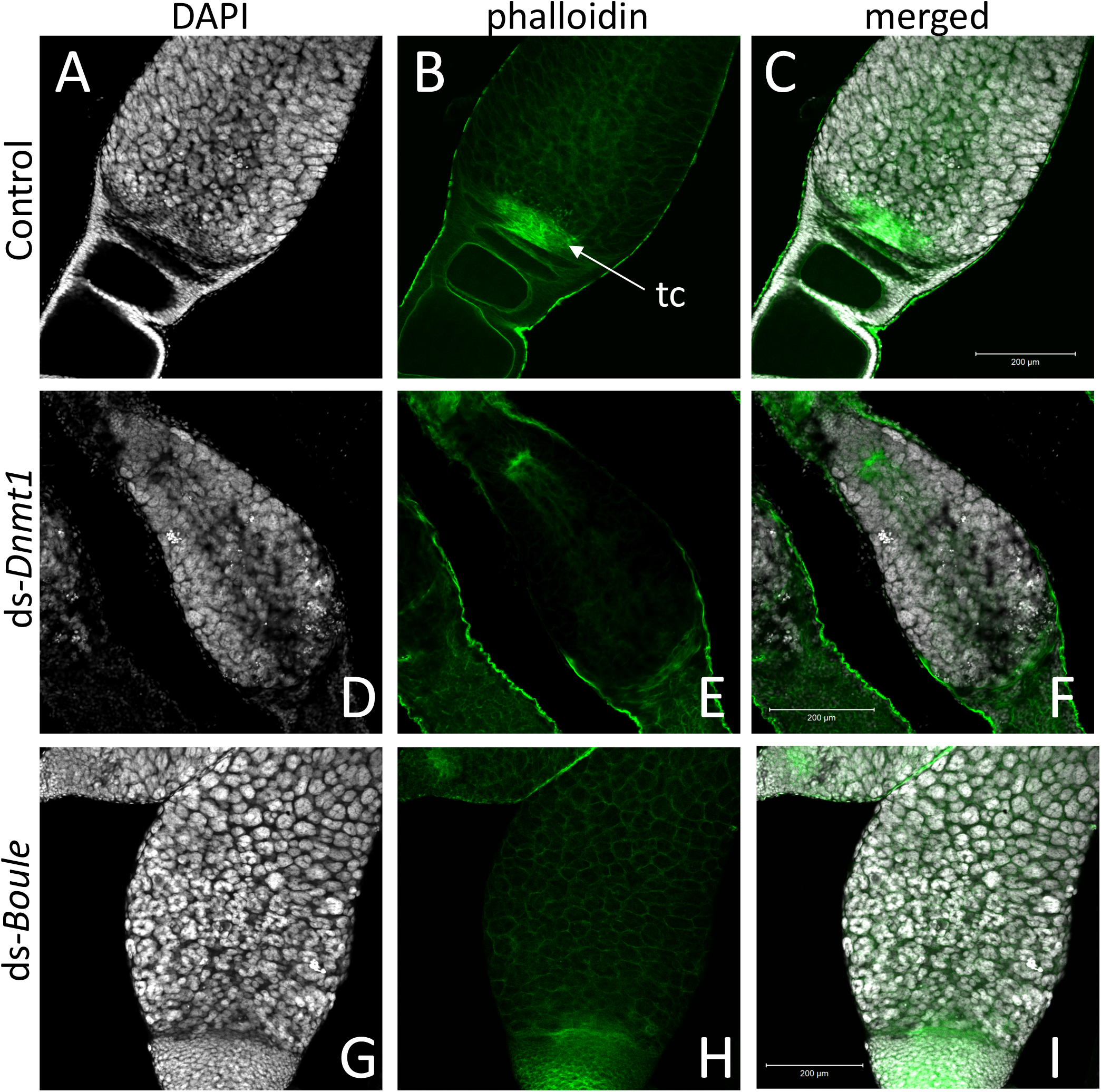
The structure of the germarium depends on the gene knockdown. In the control ovary (A-C), the large, aggregate nuclei of the trophocytes (A) were readily apparent, as are the smaller nuclei at the base of the germarium, consisting of the oogonia, primary ooctyes, and prefollicular cells. Actin staining (B) reveals both the early stages of oocyte development as the young oocytes become enveloped by follicle cells and lined by actin (B). The other structures revealed by actin staining are the developing trophic cords at the base of the germarium (tc), and the increased actin concentration within the trophic core. In ovaries from *Dnmt1* knockdown females (D-F), the trophic cells were apparent (D), as was the trophic core (E). However, there were few small nuclei at the base of the germarium, and there often showed evidence of cell death. Ovaries from *Boule* knockdown females (G-H) contained both the large trophic nuclei (G) and smaller nuclei associated with the oogonia and pre-follicular cells. However, there was no evidence of developing oocytes or trophic cords forming in either the *Dnmt1* females or *Boule* females. All images taken at 20X with 0.7 optical zoom.

The other cell type located in this region is the pre-follicular cells. To determine if these cells were also affected by *Dnmt1* knockdown, we examined cell division patterns by labeling the ovarioles with anti-phosphohistone H3 antibody, which labels dividing cells. The pre-follicular cells divide to provide follicle cells that will envelope the developing oocyte as it moved down through the ovariole. In our control females, division of the pre-follicle cells was apparent at the junction between the trophocytes and developing ovaries (Figure 7A). In the Boule knockdown females, the pre-follicular cells were particularly clear as they were not obscured by the developing ooctyes (Figure 7C). Although the germarium tissue disruption in the *Dnmt1* knockdown made it difficult to identify the pre-follicular cells structurally, there was clear evidence that they were present based on the presence of dividing cells within this region of the germarium (Figure 7B).

**Figure 7.**
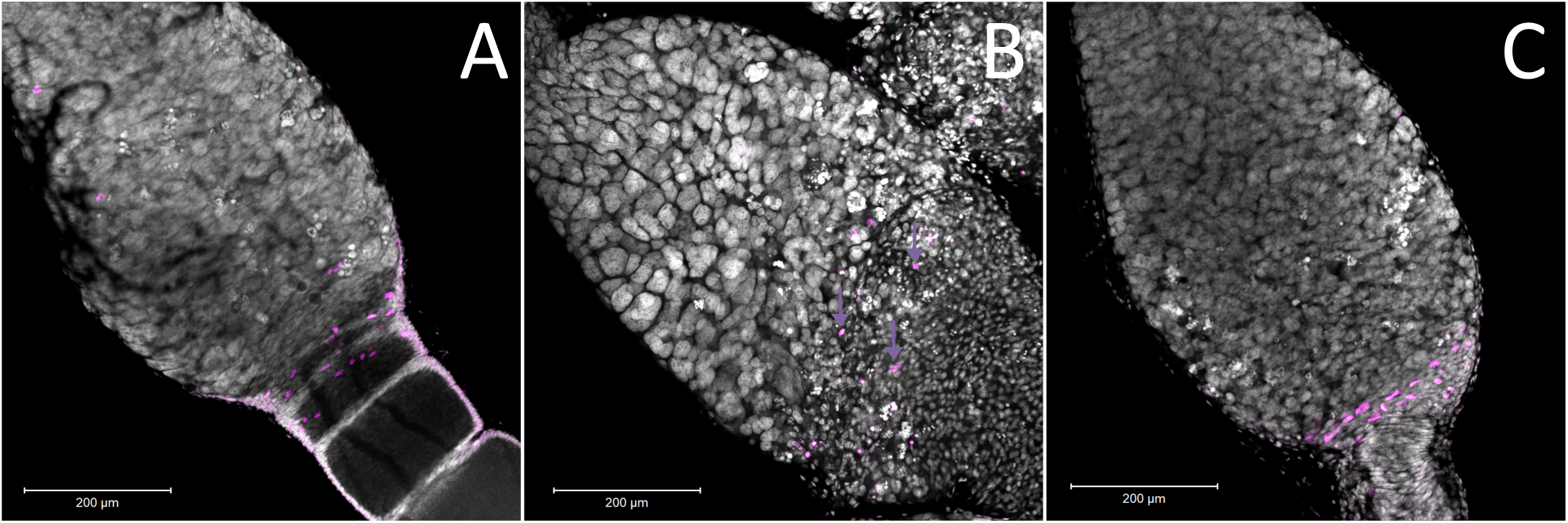
Ovaries stained for the M-phase of the cell cycle suggested that there are pre-follicular cells within all three treatments. (A) Within the control ovary, the pre-follicular cells are present in the tissues that surrounded the ooctyes at the early stages of development and the majority of the nuclei that stained positive for pHH3 were present within this region of the germarium. In the *Boule* knockdown ovaries (C), the prefollicular cells were very apparent as they were not obscured by any developing ooctyes. While the structure of the germarium in the *Dnmt1* knockdown females made it difficult to identify particular cell types, there was evidence that prefollicular cells remained in these germarium, as there are nuclei of the right size at the base end of the germarium and these were frequently stained for cell division (yellow arrows), although they were somewhat obscured by the condensed nuclei within this region of the germarium. All images were stained with DAPI and anti-phosphohistone H3 antibody and imaged at 20X with 0.7 optical zoom.

**Figure 8.**
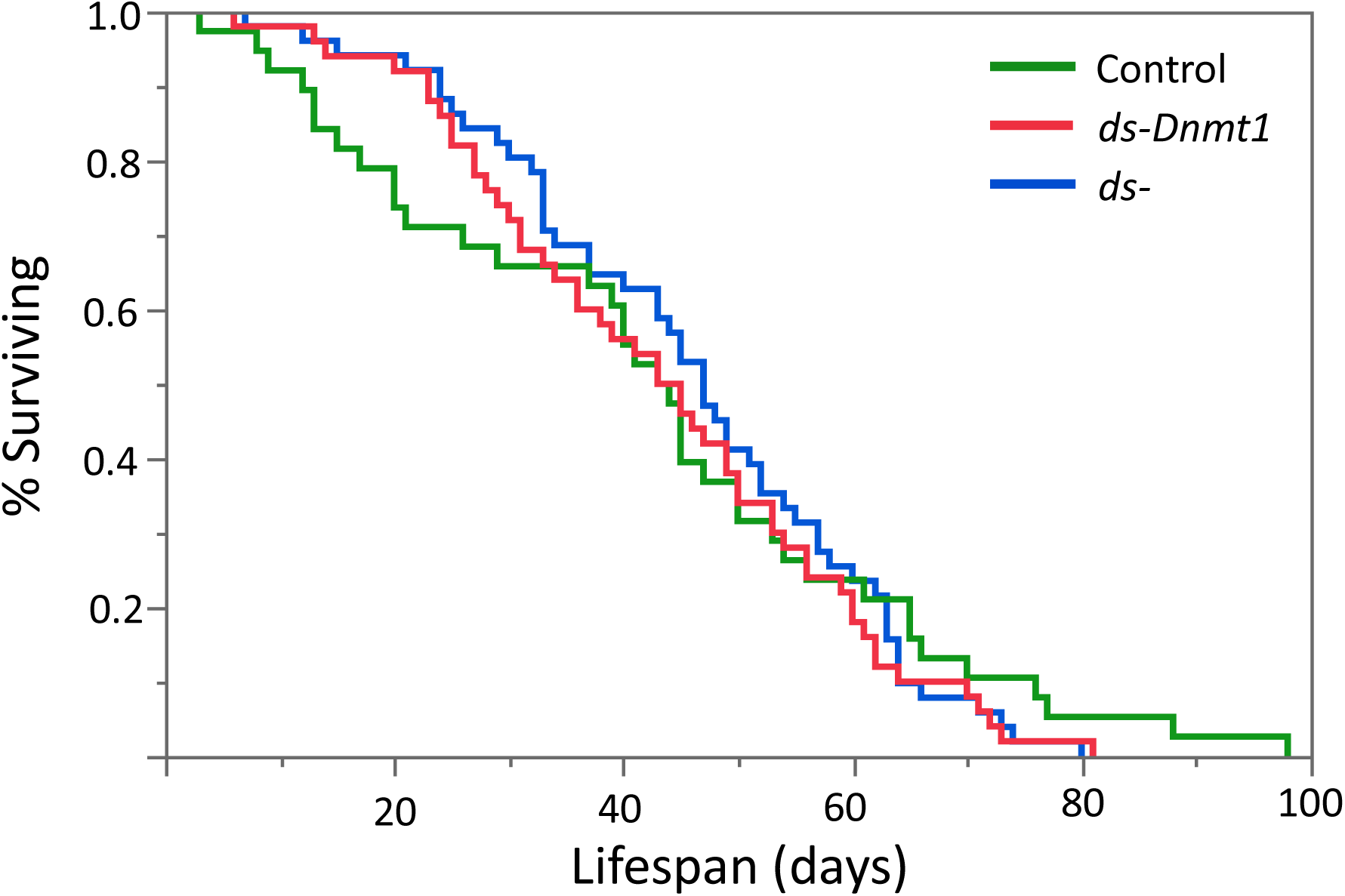
Neither the loss of egg production via *Boule* knockdown nor loss of egg production combined with loss of DNA methylation by *Dnmt1* knockdown affected female lifespan. All females were treated during the 4^th^ instar larval stage and housed individually with a male from the day of adult emergence. Lifespan was measured as the number of days between adult emergence and death.

### *Dnmt1* knockdown does not significantly affect lifespan

We predicted that if *Dnmt1* plays a role in somatic function, we would see reduction in lifespan in *Dnmt1* downregulated females due to the reduction of *Dnmt1* and DNA methylation observed in the somatic tissues as indicated by our results with gut and muscle tissue. To control for any impact on lifespan due to reduced reproductive effort, we included ds-*Boule* treated females in our lifespan analysis. We found no statistically significant differences in lifespan among the three treatments (Figure 7; Wilcoxon *χ*^2^ = 0.882, d.f. = 2, p = 0.643), indicating that somatic tissues were not negatively impacted by reduction of DNA methylation or *Dnmt1* expression.

## Discussion

There has been much discussion about the role of DNA methylation and the function of the DNA methyltransferases in insects (Glastad et al. 2019, Yan et al. 2015, Lyko and Maleszka 2011, Lo et al. 2018), given the diversity in both methylation state and patterns of evolution of the DNA methyltransferases across the insect tree of life (Bewick et al. 2016). There have been fewer experimental studies. In this study, we asked whether the phenotype observed following adult injection with ds-*Dnmt1* was restricted to the ovary because this was the only tissue that had sufficient rounds of DNA replication to reduce DNA methylation in the adult tissues or if *Dnmt1* actually has an additional function in the ovaries. We found that the germ cell specific phenotype produced by *Dnmt1* knockdown is exacerbated by knocking down *Dnmt1* function at the developmental stage at which oocytes are being born (Wick and Bonhag 1955). Thus, our results support a pleiotropic function for DNMT1, functioning as an ovary-specific gene regulating development of eggs through either the production or maintenance of oogonia as well as in the maintenance of methylation.

As evidence from more insect systems emerge, it is clear that DNMT1 is required to restore methylation following semi-conservative replication in the insects. Knocking down *Dnmt1* expression results in a reduction of DNA methylation (Bewick et al. 2019). Further, DNMT1 is required for embryo development in some insect species. Expression of *Dnmt1* is high in eggs and embryos relative to other tissues in several insect species (Yang et al. 2014, Shulz et al. 2018, this study). Knocking down *Dnmt1* expression in eggs and embryos results in embryo lethality (Zwier et al. 2012, Schulz et al. 2018, Bewick et al. 2019). Third, there is a suggestion that DNA methylation may be important in stability of germ cells (Bewick et al. 2019, Takashima et al. 2009). In honey bees, DNA methylation is more efficiently maintained in the germline than in the soma (Harris et al. 2019). This could indicate that preservation of germline DNA methylation patterns may be more critical than those of somatic methylation patterns, although it could also be a reflection of other mechanisms that stabilize the genome of the germline (Maklakov and Immler 2016). Knockdown of the DNMT1 associated protein 1 is also associated with a breakdown in ovaries and a loss of viable oocytes (Gegner et al. 2019). And loss of *Dnmt1* function leads to loss of the ability to produce ooctyes (Bewick et al. 2019).

Much of what we know about DNA methylation in insects comes from correlational studies, based on expression patterns of the DNA methyltransferases. The few studies that explicitly disrupt expression of *Dnmt1* assume that the functions revealed by this manipulation are mediated by the resulting change in DNA methylation patterns. Is DNMT1 function always related to DNA methylation? In Bewick et al. (2019), no phenotype is apparent within the somatic tissues such as the gut and muscle. However, rates of cell division in adult insects tend to be low and so the somatic tissues assayed did not lose DNA methylation. Thus, the lack of phenotype could have been due to the experimental design in which the timing of knockdown relative to assay was insufficient to allow any effect to be manifest. This study was specifically designed to address this issue; we treated nymphs to allow sufficient rounds of cell division to enable reduction of DNA methylation in both reproductive tissues and somatic tissues. We did indeed observe that the levels of DNA methylation were reduced over 2-fold in all the tissues tested. However, despite this over 2-fold reduction in DNA methylation in somatic tissues, we saw no disruption in the function of the somatic tissues. An altered function should have resulted in an impact on development and/or lifespan. Thus, DNA methylation and phenotype were uncoupled in this experiment. This is consistent with similar results in other insects. *Tribolium castaneum* is an insect with no detectable DNA methylation (Zemach et al. 2010, Cunningham, et al. 2015, Bewick et al. 2016, Schulz et al. 2018), but which has maintained the *Dnmt1* gene in its genome (Bewick et al. 2016). When *Dnmt1* is knocked down during pupal development, it results in embryo lethality (Schulz et al. 2018). The data showing that DNA methylation can be lost without a phenotype and that DNMT1 plays a functional role in an insect that lacks DNA methylation make it likely that the functional role of DNMT1 is not always mediated through DNA methylation.

Our results establish that *Dnmt1* has a specific function in germ cells. The comparison with the *Boule* knockdown females indicates that *Dnmt1* is required for maintenance of oogonia or primary oocytes. In the *Boule* knockdown females, the overall structure of the germarium is preserved, although no ooctyes ever form. Given that *Boule* is required for entry into meiosis (Eberhart et al. 1996), one likely interpretation of our data is that healthy oogonia are waiting for the signal that it is time to divide meiotically to form primary ooctyes, but that signal never comes. The tissue disruption and evidence of cell death in the *Dnmt1* knockdown females, however, indicate that *Dnmt1* is required to maintain oogonia and/or primary ooctyes and that if *Dnmt1* function is missing, these cells are not viable. While we are not able to differentiate among oogonia and primary ooctyes in our samples, the fact that the trophic tissue develops normally provides a clue as to where *Dnmt1* is required. During the first, second, and third instar stages of development, the germ cells within the somatic ovary increase in number through mitosis (Wick and Bonhag). In the fourth instar stage, oogonia either divide mitotically to form trophocytes or divide meiotically to form ooctyes.

We suggest that *Dnmt1* may be required for proper progression through meiosis or for stability of the germ cells. In *Dnmt1* knockdown females, oogonia that have not been able to advance properly through meiosis will die or be targeted for destruction. Alternatively, oogonia that divide to form primary ooctyes may require *Dnmt1* to maintain genome integrity. In the absence of *Dnmt1* newly born oocytes may degenerate. We currently are not able to determine if this is due to the loss of methylation or a pleiotropic function of DNMT1. While it is clear that DNA methyltransferases are associated with the reproductive cells in insects, the role of DNA methylation in germ cell development in male mice has been more closely examined. DNA methyltransferase 3-like (*Dnmt3L*) is required for meiosis in male mice (Bourc’his and Bestor 2004). *Dnmt3L* is required in cells prior to meiosis, perhaps involving a premeiotic genome scan that occurs in prospermatagonial cells, and the authors conclude that the defect that arises through the knockdown of expression of *Dnmt3L* is likely to arise because normal methylation patterns on dispersed repeated sequences are not properly established (Bourc’his and Bestor 2004). Knockdown of *Dnmt1* expression in male mice also impacts male fertility. Loss of *Dnmt1* expression induces apoptosis in germline stem cells, without evidence of aberrant gene expression (Takashima et al. 2009). It is possible that this is due to hypomethylation, but the authors point out that the few rounds of cell division between knockdown and sampling did not provide an opportunity for significant loss of CpG methylation. They thus conclude that mouse germline stem cells are either extremely sensitive to slight demethylation or require DNMT1 activity outside of its function in maintaining methylation state.

Given the extreme impact on fitness of *Dnmt1* in *O. fasciatus, N. vitripennis*, and *T. castaneum*, it is curious that this gene shows such variation across the insect tree of life where it is duplicated or lost across multiple taxa (Bewick et al. 2016). We expect genes with strong fitness effects, including genes essential in development, to experience strong stabilizing selection (Charlesworth 1991, Ellegren and Parsch 2007). For example, *Boule*, which is required for entry into meiosis, is a highly conserved gene across the bilaterian and experiences strong purifying selection (Shah et al. 2010). If *Dnmt1* is required for oogenesis and early development in phylogenetically disparate species, how is so much variation in this gene tolerated? Variable DNA methylation within somatic cells is typical within insects, suggesting this is not an essential function. In honey bees, levels of methylation within somatic cells vary and only have to be maintained above a threshold level within exons (Harris et al. 2019). We found a similar tolerance for changes in somatic cell levels of DNA methylation here, where an over 2-fold change in CpG methylation had no discernable effect on somatic function. It is likely, however, that there is less flexibility in the pathways required for maintaining a stable germline (Maklakov and Immler 2016), which is expected to be strongly related to fitness and therefore under strong stabilizing selection. If *Dnmt1* has a pleiotropic function in germline stability, independent of DNA methylation, it suggests that alternative or multiple pathways for stabilization of the germline genome must exist. Those pathways are yet to be discovered but examining insects that lack *Dnmt1* (Bewick et al. 2018) provides a potential model.

## Conclusion

We have demonstrated that *Dnmt1* functions to maintain a healthy germline in female *O. fasciatus. Dnmt1* knockdown, while resulting in a reduction in DNA methylation across both the germ and soma, only has a morphological phenotype in reproductive cells. We propose that *Dnmt1* has a pleiotropic function independent of DNA methylation in the germ cells and could be required to maintain the genome integrity of germ cells or be required to progress through meiosis. These results raise a number of questions that need to be addressed in future experiments.

## Data Availability Statement

The datasets analyzed for this study will be made available by the authors through the publicly available Dryad Digital Repository (https://datadryad.org/stash/)

## Author Contributions

PJM, AJM, RJS contributed conception and design of the study. AUA, JW, ECM, and PJM collected the data. All authors contributed to the analysis of the data. AUA wrote the first draft of the manuscript. All authors contributed to manuscript revision and all authors read and approved the submitted version.

## Acknowledgements

The authors thank Aleksandar Popadić and Lisa Hanna for helpful conversations about the injection protocol for larval stages of *O. fasciatus* and Christina Ethridge for preparing and analyzing low throughput whole genome bisulfite sequencing data.

