## Supplemental Table 1 and Figure 1 for "More than DNA methylation: does pleiotropy drive the complex pattern of evolution of *Dnmt1*?"

### Supplementary Materials

**Supplementary materials Table 1**.

| Treatment | Tissue | Sample Name | Raw reads | Alignment rate | Aligned reads | Conversion Rate | # methylated CpG Sites | Total CpG Sites | Percent CpG Methylation |
|---|---|---|---|---|---|---|---|---|---|
| control | gut | C_G7 | 185600 | 0.46 | 85190 | 99.84% | 14535 | 115553 | 12.57864357 |
| control | gut | C_G8 | 470549 | 0.19 | 90816 | 99.68% | 14702 | 113321 | 12.97376479 |
| control | gut | C_G9 | 250757 | 0.41 | 103989 | 99.52% | 17618 | 139298 | 12.64770492 |
| control | muscle | C_M13 | 371589 | 0.23 | 83905 | 99.75% | 14278 | 111145 | 12.84628188 |
| control | muscle | C_M14 | 172453 | 0.49 | 85123 | 99.82% | 14491 | 114420 | 12.66474393 |
| control | muscle | C_M15 | 295028 | 0.32 | 93966 | 99.19% | 16503 | 126071 | 13.0902428 |
| control | ovary | C_OV1 | 217166 | 0.42 | 91166 | 99.26% | 14659 | 118965 | 12.32211155 |
| control | ovary | C_OV2 | 202742 | 0.43 | 86976 | 99.27% | 14268 | 116081 | 12.2914172 |
| control | ovary | C_OV3 | 251823 | 0.34 | 84738 | 99.51% | 13193 | 112148 | 11.76391911 |
| ds-Dnmt1 | gut | D_G10 | 205790 | 0.47 | 96001 | 99.63% | 7618 | 129929 | 5.86320221 |
| ds-Dnmt2 | gut | D_G11 | 275442 | 0.32 | 86764 | 99.68% | 6374 | 116727 | 5.460604659 |
| ds-Dnmt3 | gut | D_G12 | 549277 | 0.18 | 98430 | 99.42% | 7730 | 126607 | 6.105507594 |
| ds-Dnmt4 | muscle | D_M16 | 364199 | 0.25 | 91887 | 99.20% | 6066 | 117124 | 5.179126396 |
| ds-Dnmt5 | muscle | D_M17 | 210362 | 0.43 | 90750 | 99.72% | 7014 | 121124 | 5.790759882 |
| ds-Dnmt6 | muscle | D_M18 | 208208 | 0.48 | 99690 | 99.82% | 5765 | 132606 | 4.347465424 |
| ds-Dnmt7 | ovary | D_Ov4 | 178014 | 0.49 | 88046 | 99.75% | 3871 | 120255 | 3.218992973 |
| ds-Dnmt8 | ovary | D_OV5 | 267613 | 0.34 | 92086 | 99.49% | 4010 | 122571 | 3.271573211 |
| ds-Dnmt9 | ovary | D_OV6 | 266599 | 0.36 | 95256 | 99.72% | 3669 | 129499 | 2.833226511 |

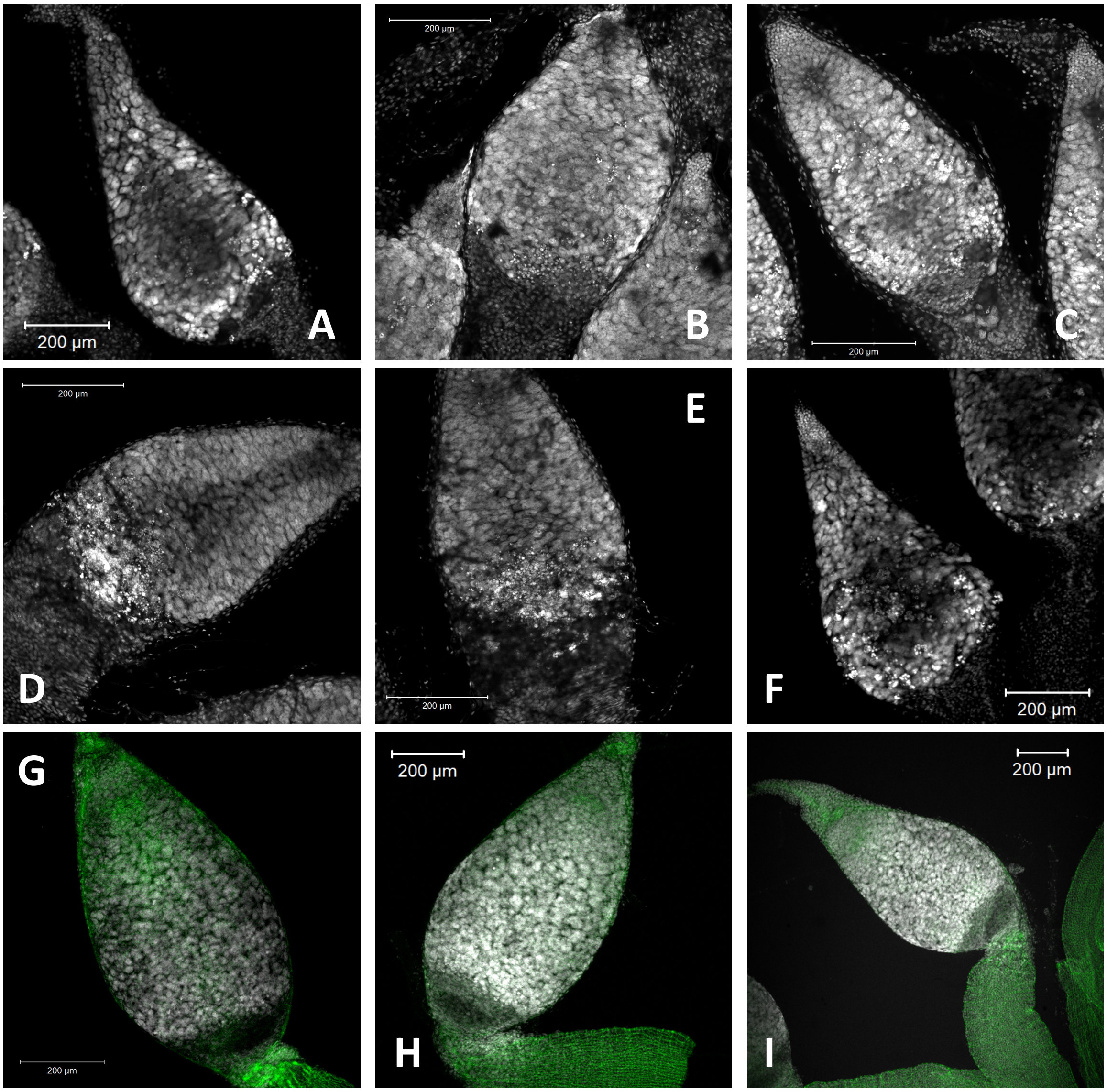


**Supplementary materials Figure S1**. A selection of images jof germaria from *Dnmt1* knockdown females showed the variety of different phenotypes observed. Many of the ovarioles contained condensed, degenerating nuclei, evidenced by DAPI staining (A-F). These degenerating nuclei were primarily located within the region of the germarium below the trophic cells, which typically presented with a normal nuclear structure. Another phenotype, most obvious in ovarioles stained with DAPI and phalloidin, that was sometimes observed was a breakdown in tissue integrity at the junction between the germarium and pedicel of the ovariole (G-H). This presented a empty spaces within the germarium. Occasionally these looked like early stage ooctyes, but is was apparent by the absence of the actin lining and any follicle cells that these are holes in the tissue and were not developing ooctyes. All images are either 10X or 20X with 0.6 optical zoom. Scale bars are all 200 µm in length.
